# Trait-Based Selection of Annual Plant Species for Enhancing Ecosystem Functioning in Extensive Green Roof Systems: A Systematic Review

**DOI:** 10.64898/2026.10.06.757100

**Authors:** Natalia Malcherczyk-Bies, Edyta Sierka

## Abstract

Extensive green roofs are an important component of urban green infrastructure. However, harsh rooftop conditions, including shallow substrates, high temperatures and limited water availability, considerably restrict plant selection. Although extensive green roofs are traditionally dominated by succulent species, increasing attention has been directed towards annual plants because of their rapid establishment, developmental plasticity and potential contribution to vegetation diversity. This review aimed to identify annual plant species introduced on extensive green roofs and to characterize the functional traits most frequently associated with their establishment under rooftop conditions. In addition, the review examined the geographical distribution and methodological characteristics of existing studies and identified species with potential for application under temperate climatic conditions. A total of 22 studies published between 2015 and 2025 were analyzed, comprising 100 annual species representing 29 plant families. The reviewed studies showed a clear geographical bias towards Mediterranean and arid regions, whereas temperate climates were considerably underrepresented. Among species occurring in Central Europe, ruderal life strategies, drought avoidance, efficient dispersal mechanisms, low plant height and prolonged flowering were the most frequently observed functional characteristics, while seed mass and specific leaf area (SLA) exhibited substantial interspecific variation. These findings suggest that functional traits, rather than taxonomic affiliation alone, provide a useful framework for selecting annual species for extensive green roofs. The review also highlights the need for long-term studies conducted under temperate climatic conditions to improve evidence-based plant selection and support the development of resilient and multifunctional green roof vegetation.

## 1. Introduction

Rising living standards and rapid economic growth have led to large-scale urban development at the expense of green areas. The lack of greenery in cities, as well as the resulting impermeable surfaces, has serious ecological consequences, such as increased temperature, concentration of air pollutants, storms, decreased air humidity and serious health problems (de Lima et al., 2023; Gutierrez et al., 2024; Santamouris and Osmond, 2020; Vijayaraghavan, 2016; Wolch et al., 2014). In response to these adverse phenomena, sustainable solutions are being implemented (Berardi et al., 2014), including the development of green infrastructure. Green roofs, or roofs with vegetation, are considered green infrastructure and often provide, at least partially, a compensation for the lack of green space in urban environments.

Along with other green elements, they are regarded as an essential component for the functioning of urban ecosystems, providing numerous benefits or ecosystem services such as contributing to temperature regulation, storm-water management, energy conservation, reduction of air pollution, and offering habitat and food for other living organisms (Francis and Jensen, 2017; Gutierrez et al., 2024; Lepczyk et al., 2017; Oberndorfer et al., 2007; Robbiati et al., 2022).

Green roofs are generally categorised as either ‘extensive’ or ‘intensive’. In urban environments, extensive green roofs (EGRs) are most implemented due to their relatively simple design, lighter structural load—resulting from a substrate layer typically no deeper than 20 cm—and lower maintenance requirements, whereas intensive green roofs necessitate a substantially thicker substrate layer, exceeding 20 cm (Oberndorfer et al., 2007).

Due to the challenging environmental conditions on green roofs (Casalini et al., 2017; Saraeian et al., 2022), especially during periods of high temperatures and limited water availability, plant species with high tolerance to environmental stress are introduced. This tolerance allows them to survive and thrive in unfavourable conditions while providing essential ecosystem services (Lundholm, 2015b). Succulents are most commonly used for extensive roof plantings because they are drought-tolerant, easy to propagate, have shallow root systems, grow low to the ground, and require minimal water and nutrients (Di Micelli et al., 2022; Lonnqvist et al., 2023; Young et al., 2017).

However, succulents do not always provide all the benefits expected from extensive green roofs (EGRs), as systems composed solely of these species are characterized by lower biodiversity and limited ecosystem services, particularly in terms of aesthetic value. Consequently, there is considerable interest in exploring alternative plant groups, with a particular focus on native species adapted to water stress (Lonnqvist et al., 2023; Saraeian et al., 2022; Zanin and Bortolini, 2020).

In this context, the integration of annual plant species into green roof design should be considered a critical step toward overcoming current limitations of conventional systems, as these species have the potential to substantially enhance functional diversity, improve temporal dynamics of vegetation, and increase the overall delivery of ecosystem services under highly variable and stress prone rooftop conditions.

Selecting suitable plant species for green roofs is most often done according to a process known as the habitat template approach (Lundholm, 2006, 2010, 2015), which suggests that plant species suitable for green roofs can be found in microclimates or habitats with biotic and abiotic conditions, such as those found on extensive green roofs. Species combinations, rather than monocultures, adapt better to limited resource availability under drought conditions and can employ different growth and survival strategies. Also, diverse species mixtures can positively affect ecosystem functioning. (Lonnqvist et al., 2023; Matsuoka et al., 2020). To improve biodiversity and the aesthetic value of urban areas, flowering perennial species are most introduced on green roofs (Caceres et al., 2022; Caceres et al., 2024; Kucuck and Eksi, 2023; Matsuoka et al., 2020). However, few studies have focused on the introduction of annual species on green roofs.

As annual plants, their life cycle is limited to a single growing season, which ends with seed production (Friedman, 2020; Hodgson et al., 2020). It means, seeds are exposed from the beginning to various environmental conditions which promotes their better adaptation and increases their chances of survival. What is more, annuals provide pollen and nectar for pollinators and serve as a visual attraction on extensive green roofs (Erickson et al., 2019; Dunnett, 2015). Annual plants also provide quick ground cover and prevent weed invasion. Moreover, in the absence of regular maintenance of green roof planted vegetations, and due to drought, the initially planted vegetation often shifts toward annual or ruderal species (Catalano et al., 2016; Schrieke and Farrell, 2021; Thuring and Dunnett, 2019) Several studies have shown that green roofs are frequently colonized by a wide range of spontaneously appearing species, with annuals being particularly prevalent due to their effective dispersal strategies and rapid life cycles (Agra et al., 2021; Aloisio et al., 2020; Bevilacqua et al., 2015). Annual species can persist on green roofs for multiple generations (Vanstockem et al., 2018) due to the formation of a soil seed bank, where seeds can remain dormant for many years, waiting for favorable conditions to germinate (Bates et al., 2015; Lonnqvist et al., 2023; Vidaller et al., 2023).

This review aims to identify annual plant species introduced on extensive green roofs and to characterize the functional traits most frequently associated with their establishment under rooftop conditions. The review further examined taxonomic patterns, the geographical distribution and methodological characteristics of existing studies, and identified species with potential for application under temperate climatic conditions, particularly in Central Europe.

## 2. Materials and Methods

### 2.1. Search strategy and study selection

The literature review was conducted between December 2025 and January 2026 using three bibliographic databases: SCOPUS, ScienceDirect, and Web of Science, which index peer-reviewed research in ecology, urban studies, and environmental sciences. The search was performed in English and limited to research articles published from 2015 onwards. The search query consisted of the following keywords combined using Boolean operators: “annual plant” AND “green roof”.

This search strategy was designed to capture studies that explicitly address the use of annual plant species in green roof systems. All records retrieved from the databases were initially screened based on their titles and abstracts (Figure 1). Articles were considered eligible if the abstract indicated that the study involved the introduction, establishment, or performance of annual plant species on extensive green roofs. Full-text screening was subsequently conducted to confirm eligibility. Studies were included if annual species were directly sown or planted into extensive green roof substrates, either alone or in combination with perennial species and/or *Sedum*. Studies focusing exclusively on intensive green roofs, green walls, or roofs without a planted vegetation layer were excluded. Duplicate records identified across the databases were removed before final selection.

**Figure 1.**
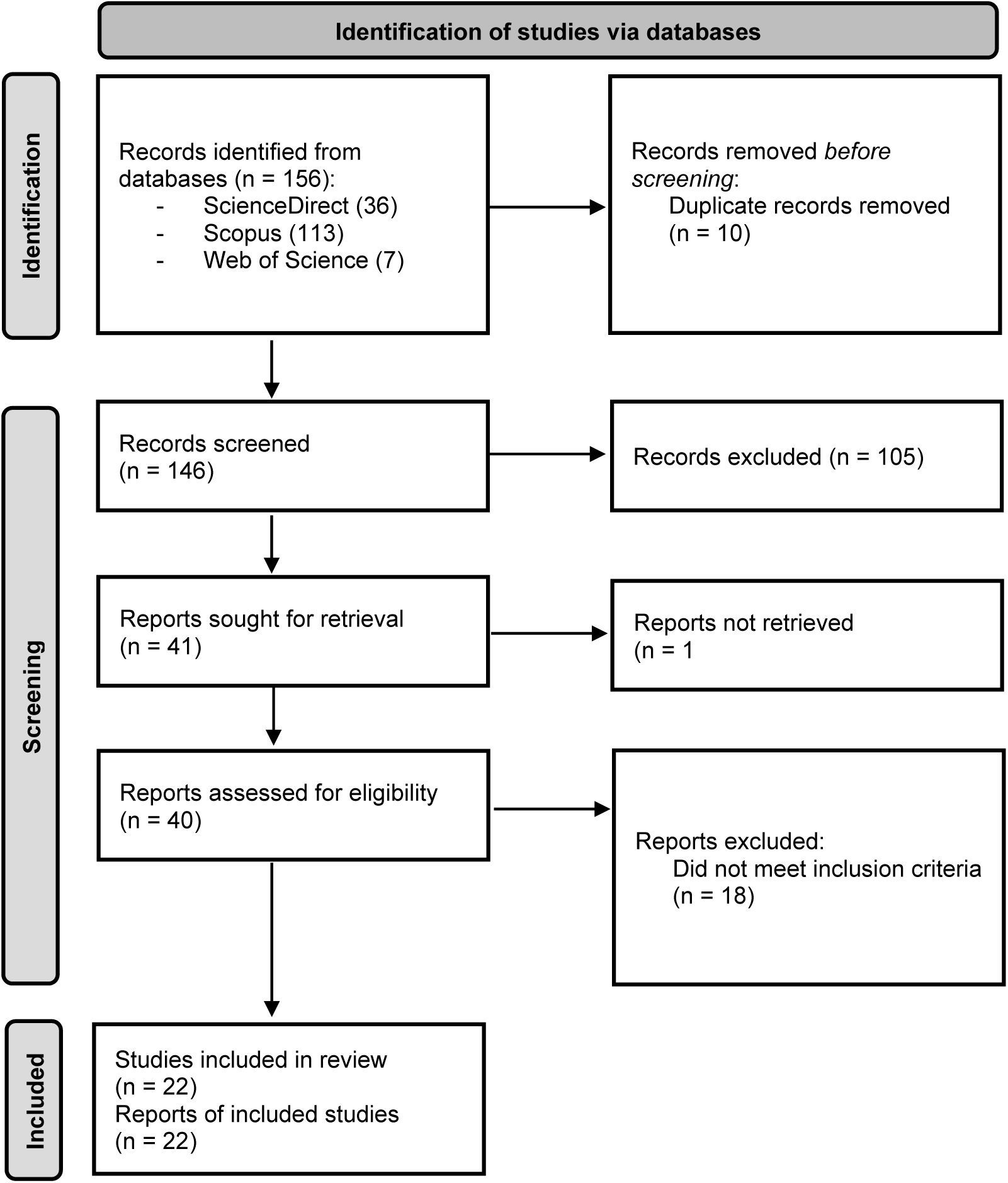
Schematic overview of the article selection process (source: Adapted from Page MJ, et al. BMJ 2021;372:n71. doi: 10.1136/bmj.n71. This work is licensed under CC BY 4.0).

The final number of articles was established by summing all selected publications from each year across the reviewed databases and subsequently eliminating duplicate records (Appendix 3).

### 2.2. Characteristic of the included studies

To provide context for the interpretation of the reported functional traits, information describing the methodological characteristics of each study was extracted. The collected data included the geographical location, climatic region, study design, experimental duration and evidence type (Table 1). These study characteristics were used to facilitate interpretation of the available evidence rather than to formally assess study quality. Because the reviewed studies differed substantially in their methodology, duration and research objectives, the evidence was synthesized qualitatively rather than quantitatively. Patterns discussed in this review were identified based on their consistency across studies employing different experimental approaches.

**Table 1.** Characteristics of the studies included in the review, including study location, climatic conditions, experimental design, and study duration.

| Study | Country | Climate zone | Study design | Duration |
| --- | --- | --- | --- | --- |
| Saraeian et al., 2025 | Australia | Temperate climate | Greenhouse experiment | 1 growing season |
| Liao et al., 2024 | Canada | Cool temperate climate | Rooftop mesocosm experiment | 1 growing season |
| Lonnqvist et al., 2023 | Sweden | Cool temperate climate | Greenhouse experiment | 1 growing season |
| Vannucchi et al., 2023 | Italy | Mediterranean | Rooftop mesocosm experiment | 7 growing seasons |
| Vidaller et al., 2023 | France | Mediterranean | Rooftop mesocosm experiment | 9 growing seasons |
| Latini et al., 2022 | Italy | Mediterranean | Green roof experiment | 3 growing seasons |
| Saraeian et al., 2022 | Australia | Temperate climate | Rooftop mesocosm experiment | 1 growing season |
| Vannucchi et al., 2022 | Italy | Mediterranean | Rooftop mesocosm experiment | 6 growing seasons |
| Schrieke and Farrell, 2021 | Australia | Temperate climate | Greenhouse experiment | 1 growing season |
| Schroder and Kiehl, 2021 | Germany | Temperate climate | mesocosm experiment | 4 growing seasons |
| Schroder and Kiehl, 2020 | Germany | Temperate climate | mesocosm experiment | 4 growing seasons |
| Matsuoka et al., 2019 | Japan | Humid subtropical climate | Greenhouse experiment | 2 growing seasons |
| Schindler et al., 2019 | Israel | Mediterranean | Rooftop mesocosm experiment | 1 growing season |
| Vasl et al., 2019 | Israel | Mediterranean | Rooftop mesocosm experiment | 2 growing seasons |
| Salman and Blaustein, 2018 | Israel | Mediterranean | Rooftop mesocosm experiment | 1 growing season |
| Schindler et al., 2018 | Israel | Mediterranean | Green roof experiment | 2 growing seasons |
| Vannucchi et al., 2018 | Italy | Mediterranean | Rooftop mesocosm experiment | 2 growing seasons |
| Agra et al., 2017 | Israel | Mediterranean | Rooftop mesocosm experiment | 4 growing seasons |
| Vasl et al., 2017 | Israel | Mediterranean | Green roof experiment | 3 growing seasons |
| Aloisio et al., 2016 | USA | Humid subtropical climate | Rooftop mesocosm experiment | 1 growing season |
| Bates et al., 2015 | UK | Temperate maritime climate | Rooftop mesocosm experiment | 6 growing seasons |
| Van Mechelen et al., 2015 | Belgium / France | Temperate maritime climate / Mediterranean | Rooftop mesocosm experiment | 3 growing seasons |

### 2.3. Species selection and trait analysis

The selection of plant species was based on predefined criteria, including their occurrence within climatic conditions comparable to those of Central Europe and the availability of functional trait data. To identify species with potential for temperate climates, the final trait analysis focused on taxa naturally occurring or widely naturalized in Central Europe. Species restricted to Mediterranean climatic conditions were therefore not included in this final subset. The selected species were subsequently characterized according to traits relevant to green roof performance, including growth form, plant height, dispersal mechanism, ecological indicator values for light and moisture, drought response, CSR strategy, flowering duration, seed mass and specific leaf area (SLA) (Table 3).

Functional trait data were obtained from the LEDA Traitbase and Pladias Traitbase, which provide comprehensive information on life-history traits of Northwest European flora (Kleyer et al., 2008; https://uol.de/en/landeco/research/leda/data-files; https://pladias.cz/en/). Predominant drought response strategies were assigned based on published ecological descriptions of each species and their functional characteristics reported in the literature.

The extracted data were synthesized using a descriptive and comparative approach to identify recurring taxonomic patterns, frequently used species, and trait combinations associated with successful establishment on extensive green roofs (Fig. 1).

## 3. Results

After a rigorous selection process, 22 articles were identified that specifically address the direct introduction of annual plant species onto the substrate of extensive green roofs. These articles were published between 2015 and 2025, with the number of publications remaining relatively consistent over this period. Of the 22 studies, only four focused exclusively on annual species, while five introduced both annuals and *Sedum* species. The remaining 13 studies included a mixture of annual and perennial species in their plantings (Fig. 2). The studies included in the selected articles were conducted across 11 countries (Table 1). Israel accounted for the highest number of studies investigating the use of annual species on green roofs. A notable level of research activity focusing on annual species was also observed in Australia and Italy.

**Figure 2.**
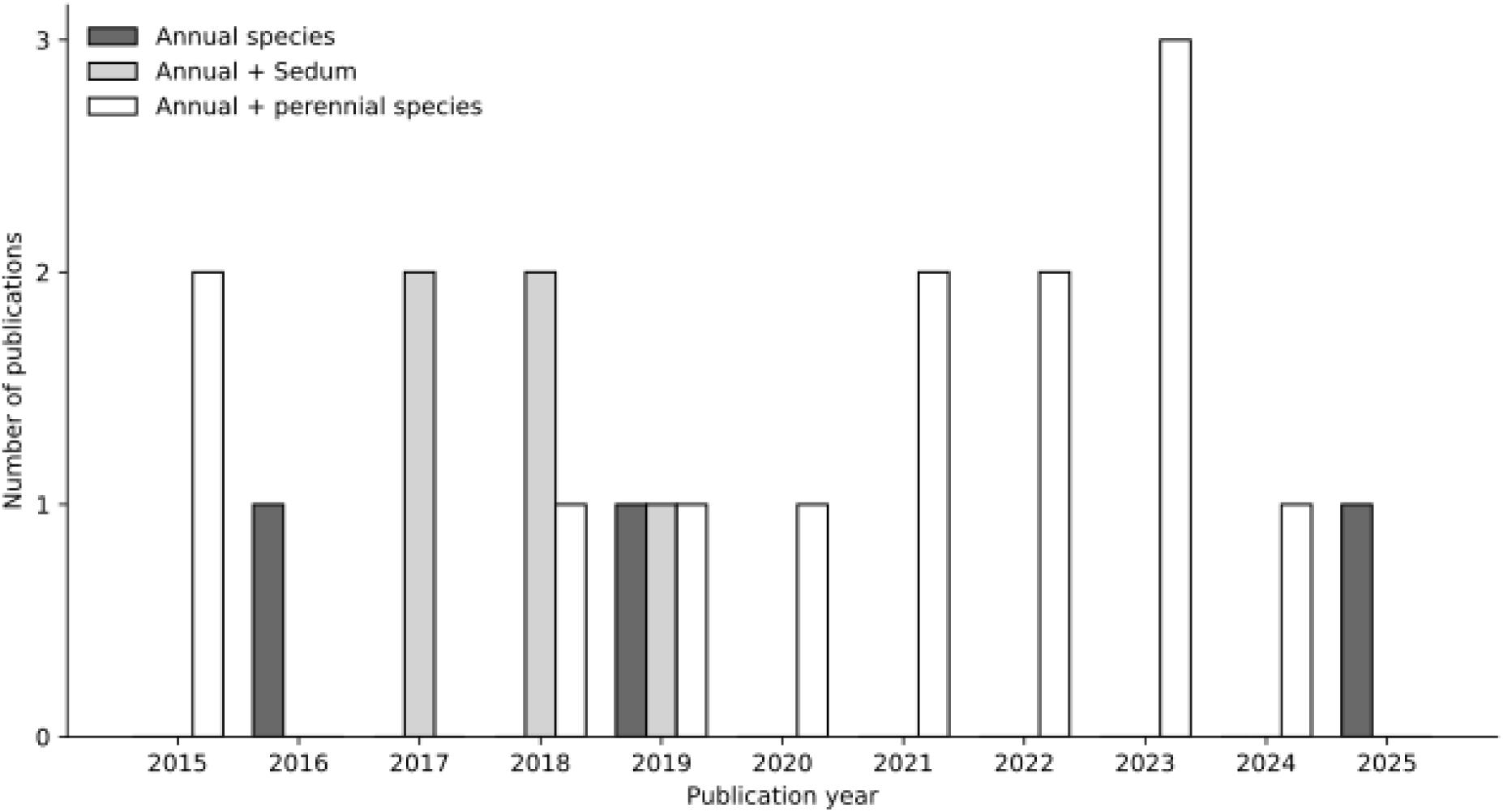
Number of publications (2015–2025) reporting the introduction of plant species onto extensive green roofs, grouped according to the vegetation type investigated: annual species, annual species combined with *Sedum*, and annual species combined with perennial species.

From the selected articles, 100 annual plant species belonging to 29 families were identified (Appendix 1). The most frequently introduced species included *Lagurus ovatus*, *Agrostemma githago*, *Alyssum alyssoides*, *Anthemis pseudocotula*, *Chrysanthemum coronarium*, *Echium judaeum*, *Ricotia lunaria*, *Silene aegyptiaca*, *Stipa capensis*, and *Trifolium arvense* (Table 2). Among all introduced species, the family Asteraceae was the most represented, with 25 species (Fig. 3).

**Figure 3.**
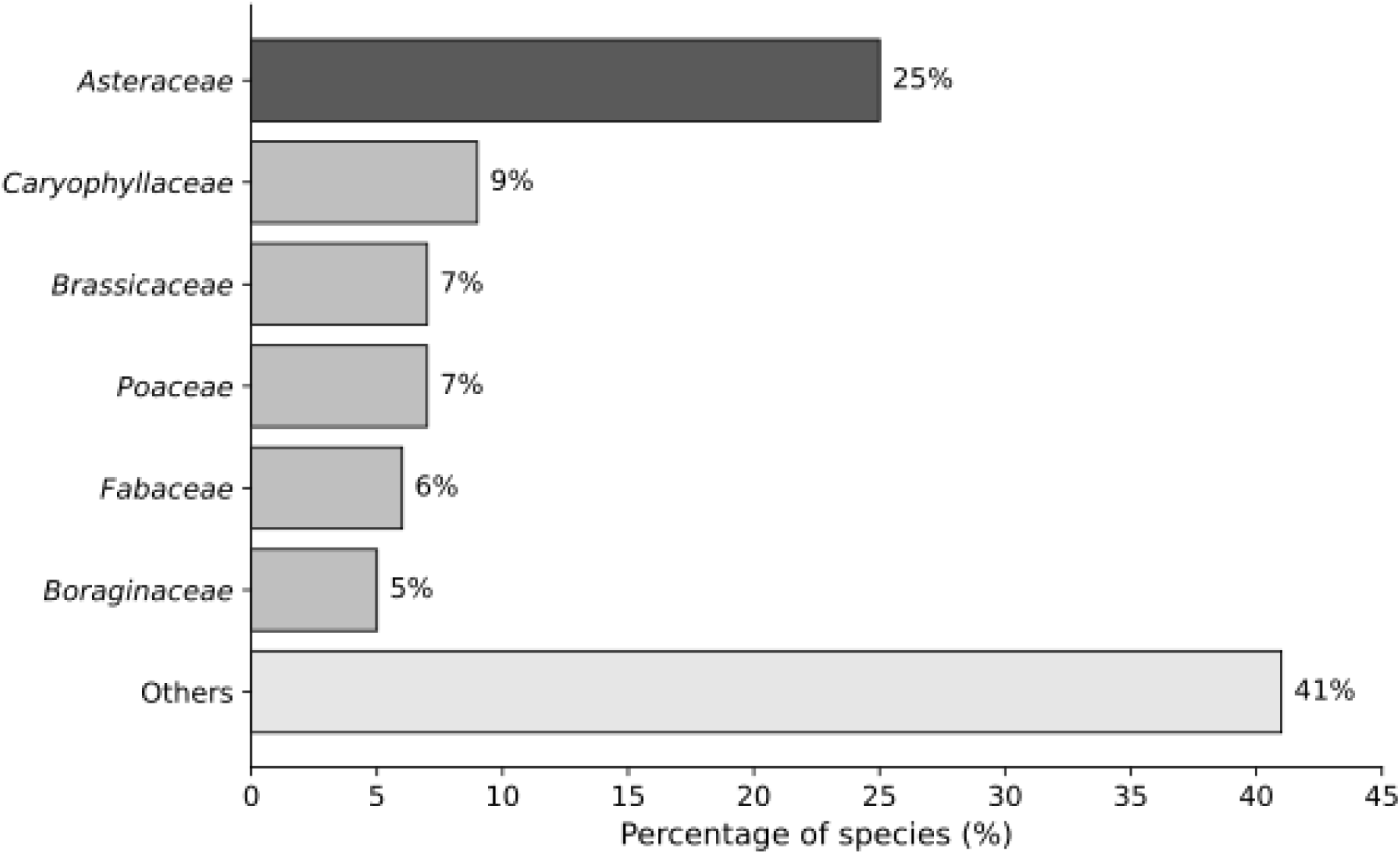
Percentage distribution of annual plant species among the most represented plant families identified in the reviewed studies. Plant families represented by fewer than five species were grouped into the category “Others”. Percentages are based on the total number of species identified across the reviewed studies (n = 100).

**Table 2.** Most frequently introduced plant species on green roofs identified in the analyzed studies. The frequency value represents the number of articles in which each species was recorded.

| Number of introduced species | 3 | 4 | 5 | 6 |
| --- | --- | --- | --- | --- |
| Species | *Calendula arvensis*<br>*Capsella bursa-pastoris*<br>*Geranium molle*<br>*Malva parviflora*<br>*Portulaca oleracea*<br>*Reseda alba*<br>*Silene gallica*<br>*Tordylium carmeli* | *Cichorium endivia*<br>*Chaetosciadium trichospermum*<br>*Erodium cicutarium*<br>*Erodium malacoides*<br>*Lomelosia prolifera*<br>*Ranunculus scandicinus*<br>*Trifolium stellatum* | *Agrostemma githago*<br>*Alyssum alyssoides*<br>*Anthemis pseudocotula*<br>*Chrysanthemum coronarium*<br>*Echium judaeum*<br>*Ricotia lunaria*<br>*Silene aegyptiaca*<br>*Stipa capensis*<br>*Trifolium arvense* | *Lagurus ovatus* |

The annual species selected for temperate climatic conditions shared several functional traits relevant to their use on extensive green roofs (Table 3, Fig. 4). Ruderal strategies predominated among the analyzed taxa, with 53.8% of the species classified as R strategists, whereas CR (23.1%), SR (15.4%), and intermediate R/SR (7.7%) strategies occurred less frequently. Drought avoidance was the most common drought-response strategy (69.2%), followed by drought escape (46.2%), while physiological drought tolerance was recorded in 30.8% of the analyzed species. The frequency of the predominant functional traits is presented in Fig. 4. Most species exhibited an erect growth form (76.9%), whereas prostrate growth forms were observed in only 7.7% of the analyzed taxa. Plant height ranged from 0.25 to 1.00 m (mean = 0.48 m; median = 0.40 m), with most species not exceeding 0.5 m. Flowering duration ranged from 3 to 9 months (mean = 4.85 months). Seed mass showed considerable variation, ranging from 0.107 to 16.58 mg (median = 0.56 mg). Similarly, SLA values ranged from 17.73 to 36.50 mm² mg⁻¹ (mean = 25.79 mm² mg⁻¹).

**Figure 4.**
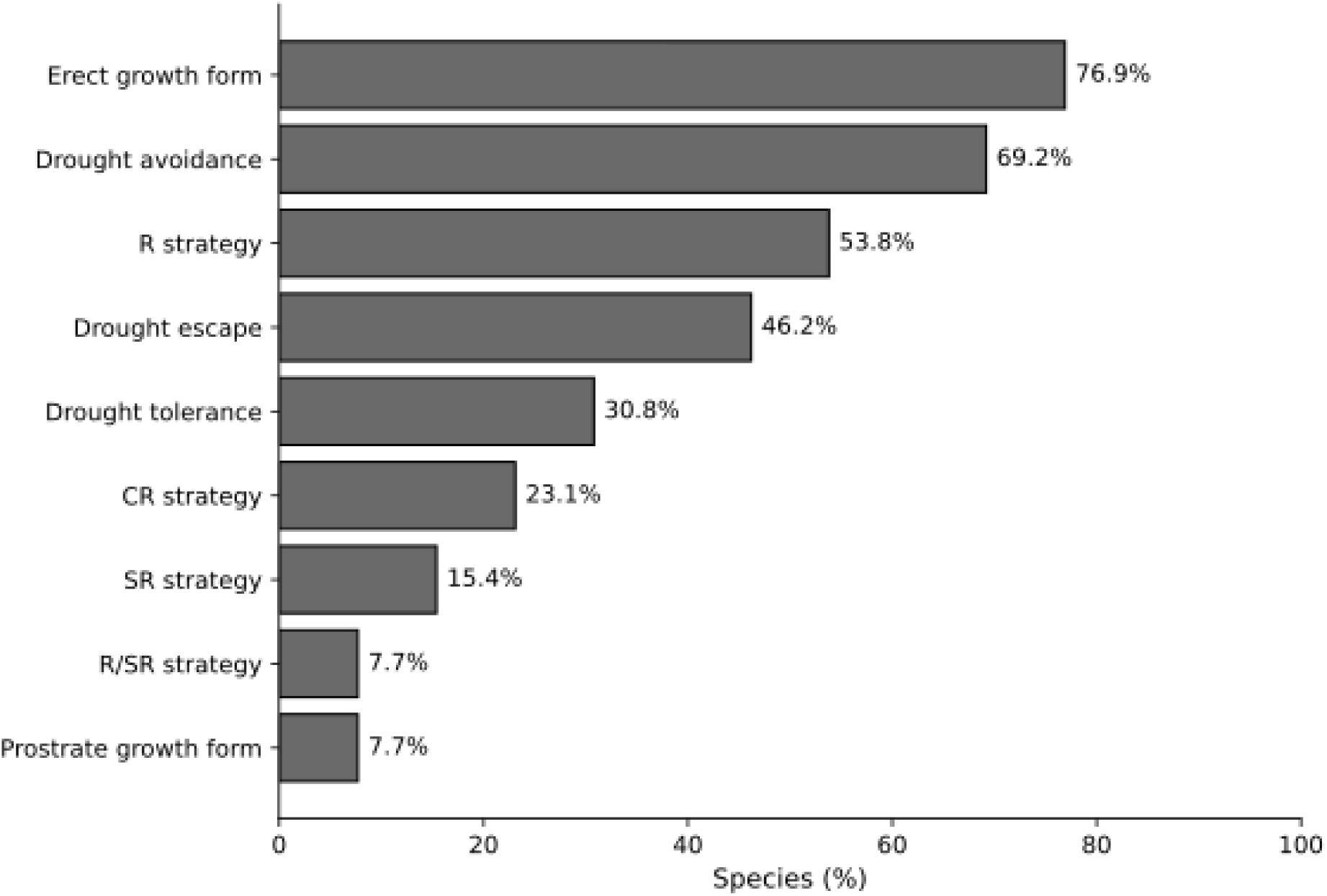
Frequency (%) of the predominant functional traits identified among annual plant species selected for extensive green roofs under temperate climatic conditions. Percentages indicate the proportion of analyzed species exhibiting each functional trait.

**Table 3.** Most frequently occurring species in extensive green roofs within the temperate climatic conditions of Central Europe and their functional traits. NA indicates that no information was available for the respective trait value.

| species | life strategy | Predominant drought response strategy | height | shoot growth form | dispersal type | seed mass [mg] | SLA [mm<sup>2</sup> / mg] | Flowering duration (months) |
| --- | --- | --- | --- | --- | --- | --- | --- | --- |
| *Lagurus ovatus* | R/SR | Avoidance + escape | 0,15 - 0,3 | stem erect | anemochory, epizoochory | 0,55 | NA | 4 |
| *Agrostemma githago* | CR | Escape | 0,3 - 1 | stem erect | autochory, barochory | 16,58 | 17,73 | 3 |
| *Alyssum alyssoides* | SR | Avoidance + escape | 0,05 - 0,3 | Stem ascending to prostrate / stem erect | autochory, anemochory | 0,415 | 18,64 | 4 |
| *Trifolium arvense* | SR | Avoidance | 0,1 - 0,3 | Stem ascending to prostrate / stem erect | epizoochory, autochory | 0,305 | 18,9 | 4 |
| *Cichorium endivia* | CR | Avoidance | 0,3 - 1 | stem erect | anemochory | 1,2 | NA | 3 |
| *Erodium cicutarium* | R | Avoidance + tolerance | 0,1 - 0,5 | stem erect | autochory, ballochory, epizoochory | 1,445 | NA | 7 |
| *Calendula arvensis* | R | Avoidance + escape | 0,1 - 0,25 | stem erect | epizoochory, zoochory, myrmecochory, anemochory | 3,66 | 29,33 | 5 |
| *Capsella bursa-pastoris* | R | Escape | 0,05 - 0,5 | stem erect | autochory, barochory | 0,107 | 32,1 | 9 |
| *Geranium molle* | R | Avoidance | 0,1 - 0,4 | stem erect | autochory, ballochory, epizoochory | 1,24 | 36,5 | 5 |
| *Malva parviflora* | R | Tolerance + avoidance | 0,15 - 0,4 | stem erect | zoochory, endozoochory, barochory | NA | NA | 6 |
| *Portulaca oleracea* | R | Tolerance | 0,05 - 0,3 | Stem prostrate | autochory, barochory | 0,45 | 18,45 | 4 |
| *Reseda alba* | CR | Tolerance + Avoidance | 0,3 – 0,8 | stem erect | autochory, myrmecochory | 0,562 | 26 | 5 |
| *Silene gallica* | R | Avoidance + escape | 0,08 - 0,25 | stem erect | autochory, anemochory | 0,384 | 34,5 | 4 |

## 4. Discussion

### 4.1. Selection annual plant species for green roofs

Drought represents one of the principal environmental constraints limiting plant establishment and persistence on extensive green roofs. Consequently, successful species are characterized not by taxonomic affiliation but by functional traits that enable them to tolerate shallow substrates, high solar radiation, fluctuating water availability and strong wind exposure. The present review supports a trait-based approach to plant selection, suggesting that combinations of functional traits rather than species identity determine adaptation to rooftop environments. Among the species selected for temperate climatic conditions, ruderal (R) strategists predominated, accounting for more than half of the analyzed taxa (53.8%), whereas intermediate CSR strategies occurred less frequently (Table 3, Fig. 4). This predominance indicates that rapid establishment, early reproduction and efficient colonization are key ecological strategies under rooftop conditions, where vegetation is regularly exposed to environmental disturbance. Similar observations were reported by Saraeian et al. (2025), who suggested that annual species possessing ruderal life strategies rapidly exploit favorable environmental conditions through accelerated growth and reproduction. Likewise, Lönnqvist et al. (2023) showed that CSR strategy is closely associated with plant water-use patterns, with ruderal species exhibiting greater phenological flexibility under changing moisture conditions. Drought-response strategy also emerged as a major determinant of species suitability. The predominance of drought avoidance (69.2%) and drought escape (46.2%) among the analyzed species indicates that annual plants primarily persist by reducing exposure to water stress through phenological adjustment rather than relying exclusively on physiological tolerance. Completing the life cycle before prolonged drought, together with developmental plasticity, enables annual species to maintain reproductive success under unpredictable water availability. Experimental studies have suggested that annual plant communities can maintain high vegetation cover and functional diversity despite different irrigation regimes, suggesting that phenological flexibility represents an effective mechanism for coping with rooftop drought (Saraeian et al., 2022). Similarly, Schrieke and Farrell (2021) reported that spontaneous annual species successfully established on extensive green roofs because they combined rapid growth with efficient regulation of water use.

Efficient dispersal was another recurring functional characteristic among the selected species. Autochory, anemochory and zoochory were the predominant dispersal mechanisms, facilitating rapid colonization of newly available rooftop habitats and vegetation recovery following disturbance. Such regeneration capacity is particularly important in extensive green roofs, where environmental conditions frequently create open microsites suitable for recolonization. Moreover, the ability of annual species to contribute to persistent soil seed banks further increases community resilience and supports spontaneous regeneration after periods of environmental stress (Vanstockem et al., 2018).

Considerable variation was observed in seed mass and specific leaf area (SLA), indicating the coexistence of contrasting ecological strategies among successful annual species. Small-seeded species generally exhibit greater dispersal capacity and colonization potential, whereas larger seeds provide greater nutrient reserves, improving seedling establishment under stressful conditions (Hodgson et al., 2020). Likewise, differences in SLA reflect variation in resource-acquisition strategies, with higher SLA generally associated with rapid growth and lower SLA indicating more conservative resource use. Rather than identifying a single optimal trait value, these findings suggest that functional diversity may increase the stability and adaptability of green roof vegetation, consistent with previous studies highlighting positive relationships between trait diversity, ecosystem functioning and vegetation persistence (Heim et al., 2023). Structural traits also appear to contribute to successful rooftop establishment. Most of the analyzed species did not exceed 0.5 m in height and were characterized by erect or creeping growth forms. Low stature probably reduces mechanical damage caused by wind while maintaining stable vegetation cover on shallow substrates. At the same time, flowering duration ranged from three to nine months, indicating substantial variation in reproductive phenology. Species with prolonged flowering periods may enhance the ecological value of green roofs by providing floral resources over an extended period, thereby supporting pollinators and increasing ecosystem multifunctionality (Erickson et al., 2019; Francis and Jensen, 2017).

Although Asteraceae was the most frequently represented family among the reviewed studies, this pattern should not be interpreted as evidence that taxonomic affiliation alone determines suitability for green roofs. Instead, many Asteraceae species combine several advantageous functional traits, including ruderal life strategies, efficient dispersal, prolonged flowering and high reproductive capacity (Waheed and Arshad, 2024; Xu and Chang, 2017). Consequently, species selection should prioritize functional trait combinations rather than family membership, as unrelated taxa possessing similar ecological characteristics may perform equally well under rooftop conditions.

### 4.2. Influence of climatic conditions on annual species selection

The geographical distribution of the reviewed studies revealed a clear predominance of research conducted in Mediterranean and arid climatic regions, particularly Italy, Israel and Australia. These regions are characterized by prolonged summer droughts, high solar radiation and elevated temperatures, making them suitable locations for evaluating plant responses to severe rooftop stress (Catalano et al., 2016; Dunnet and Kingsbury, 2004; Leotta et al., 2023). Consequently, many of the annual species investigated in these studies have been selected primarily for their ability to withstand water limitation and high thermal loads. Although Mediterranean environments provide valuable insights into drought adaptation, direct transfer of these findings to temperate regions should be approached with caution. Extensive green roofs in Central Europe experience a different combination of environmental constraints, including seasonal frost, repeated freeze–thaw cycles and greater annual variation in temperature and precipitation. Therefore, plant species suitable for Mediterranean climates may not necessarily perform equally well under temperate climatic conditions.

To address this limitation, the present review identified a subset of annual species naturally occurring or widely naturalized in Central Europe that combine functional traits associated with successful rooftop establishment. These species were characterized predominantly by ruderal life strategies, drought avoidance, efficient dispersal mechanisms and relatively low stature. Such trait combinations suggest that successful species in temperate climates must tolerate both periodic drought during summer and low winter temperatures while maintaining rapid regeneration capacity (Bellini et al., 2024; Seyedabadi et al., 2021). Similar conclusions have been reported by Schröder and Kiehl (2021), who demonstrated that annual species native to temperate regions successfully established on extensive green roofs without irrigation owing to their ecological plasticity. Likewise, Lönnqvist et al. (2023) emphasized that plant functional strategies are strongly influenced by climatic context and that responses observed under Mediterranean conditions cannot be directly extrapolated to cooler regions. These findings reinforce the need for region-specific species selection based on functional traits rather than species lists developed for other climatic zones.

The limited number of studies conducted in Central Europe also highlights an important knowledge gap. Future research should prioritize long-term field experiments evaluating annual species under temperate climatic conditions, particularly in relation to drought resilience, winter survival and interactions with perennial vegetation. Such studies would improve the transferability of trait-based recommendations and may facilitate the development of climate-adapted green roof vegetation for Central European cities.

### 4.3. Practical implications for ecological engineering

The present review provides a practical framework for the selection of annual plant species for extensive green roofs under temperate climatic conditions. Rather than relying on taxonomic composition, species selection should prioritize combinations of functional traits directly associated with successful establishment and persistence under rooftop stress (Saraeian et al., 2025; Todeschini and Fett-Neto, 2025).

The analyzed species most frequently combined ruderal life strategies, drought avoidance, efficient dispersal mechanisms, low plant height and prolonged flowering periods. These traits are particularly relevant for extensive green roofs, where vegetation must establish rapidly, regenerate after disturbance and tolerate strong fluctuations in water availability while requiring minimal maintenance. The predominance of drought avoidance and drought escape strategies suggests that annual species are especially suitable for systems exposed to periodic summer drought, as they can adjust their phenology and complete reproduction before severe water limitation occurs (Saraeian et al., 2022; 2025).

Low plant stature appears to be another important design criterion. Most of the selected species remained below 0.5 m in height, reducing the risk of wind damage and limiting structural loading while maintaining effective substrate coverage. Such characteristics are advantageous for lightweight extensive green roofs, where vegetation must remain stable under exposed urban conditions (Heim et al., 2025; Tian et al., 2019).

The observed variation in seed mass and SLA indicates that no single trait combination is universally optimal. Instead, combining species representing contrasting regeneration and resource-use strategies may increase vegetation resilience and improve long-term persistence. For example, small-seeded species may enhance colonization of newly available microsites, whereas larger-seeded species may improve establishment under stressful conditions through greater nutrient reserves (Hodgson et al., 2020; Vanstockem et al., 2018).

Extended flowering duration may further increase the ecological value of green roofs by providing floral resources over longer periods and supporting urban pollinator communities. Incorporating annual species with complementary flowering phonologies could therefore enhance the temporal continuity of ecosystem services.

Overall, the findings support a trait-based approach to green roof design consistent with the principles of ecological engineering. Selecting species according to functional traits rather than taxonomic identity may improve vegetation establishment, increase resilience to climatic

### 4.4. Limitations and future research

Despite providing a comprehensive overview of annual plant species used on extensive green roofs, this review has several limitations that should be acknowledged. First, the literature search was restricted to English language publications indexed in three scientific databases (Scopus, Web of Science and ScienceDirect). Consequently, relevant studies published in other languages or indexed elsewhere may have been overlooked. Second, the review included a relatively small number of studies (n = 22), reflecting the limited research attention devoted to annual species compared with perennial vegetation traditionally used on green roofs.

Another important limitation is the considerable heterogeneity of the analyzed studies. The reviewed publications differed in climatic conditions, experimental duration, study design and vegetation composition, ranging from controlled mesocosm experiments to long-term field studies and observations of spontaneous vegetation. As a result, the available evidence varied in its level of robustness, preventing direct quantitative comparisons among studies. For this reason, the present review adopted a descriptive trait-based synthesis rather than a formal meta-analysis.

Although the identified functional trait patterns were generally consistent across studies, the geographical distribution of available research remains strongly biased towards Mediterranean and arid regions. Consequently, knowledge regarding the performance of annual species under temperate climatic conditions, particularly in Central Europe, is still limited. Additional long-term field experiments conducted under different climatic conditions are therefore required to validate the transferability of the proposed trait combinations.

Future research should integrate functional trait analyses with quantitative approaches, including meta-analysis and multivariate methods, to identify the combinations of traits that most strongly determine plant performance on green roofs. Particular attention should be given to seed mass, specific leaf area (SLA), flowering phenology, dispersal mechanisms and drought-response strategies, as well as to interactions between annual and perennial species within mixed plant communities. Such studies would contribute to the development of evidence-based guidelines for designing resilient, multifunctional and climate-adapted green roof vegetation.

## 5. Conclusion

This review suggests that annual plant species represent an underutilized but promising component of extensive green roof vegetation. Their successful establishment is primarily determined by functional traits rather than taxonomic affiliation, supporting the application of a trait-based approach to plant selection.

Among the analyzed species, ruderal life strategies, drought avoidance, efficient dispersal mechanisms, low plant height and prolonged flowering were the most frequently occurring characteristics associated with successful establishment under rooftop conditions. Considerable variation in seed mass and specific leaf area (SLA) further indicates that combining species with contrasting regeneration and resource-use strategies may enhance the resilience and stability of green roof vegetation.

The review also revealed a strong geographical bias towards Mediterranean and arid regions, highlighting the need for further studies conducted under temperate climatic conditions, particularly in Central Europe. The identification of annual species naturally occurring in this region provides a basis for developing climate-adapted plant assemblages capable of tolerating both summer drought and winter frost.

From an ecological engineering perspective, these findings provide practical guidance for designing more resilient and multifunctional green roofs. Plant selection should prioritize combinations of functional traits that promote rapid establishment, regeneration and adaptation to environmental stress, rather than relying on taxonomic composition alone.

Future research should focus on long-term field experiments and quantitative trait-based analyses to determine which combinations of functional traits most effectively enhance vegetation persistence and ecosystem service delivery under different climatic conditions.

---

**Appendix 1.** List of introduced plant species based on selected articles, including information on their taxonomic families and natural habitats.

| Species | Family | Natural habitat |
| --- | --- | --- |
| *Agrostemma githago* | *Caryophyllaceae* | arable fields, field margins |
| *Alyssum alyssoides* | *Brassicaceae* | dry grasslands, sandy and rocky sites |
| *Amaranthus cruentus* | *Amaranthaceae* | disturbed areas, fields, roadsides |
| *Amaranthus dubius* | *Amaranthaceae* | disturbed habitats, roadsides, cultivated areas |
| *Amaranthus tricolor* | *Amaranthaceae* | fields, wastelands, disturbed sites |
| *Anthemis pseudocotula* | *Asteraceae* | fields, roadsides, wastelands |
| *Arenaria serpyllifolia* | *Caryophyllaceae* | dry soils, walls, roadsides |
| *Avena sterilis* | *Poaceae* | grasslands, fields, roadsides |
| *Brachyscome iberidifolia* | *Asteraceae* | grasslands, open woodlands, sandy soils |
| *Blackstonia perfoliata* | *Gentianaceae* | dry grasslands, coastal areas |
| *Calandrinia eremaea* | *Montiaceae* | arid areas, sandy soils, open habitats |
| *Calandrinia flava* | *Montiaceae* | dry habitats, sandy soils, shrublands |
| *Calendula arvensis* | *Asteraceae* | fields, grasslands, roadsides |
| *Capsella bursa-pastoris* | *Brassicaceae* | fields, roadsides, disturbed habitats |
| *Centaurea cyanus* | *Asteraceae* | arable fields, grasslands |
| *Centranthus macrosiphon* | *Caprifoliaceae* | rocky slopes, dry grasslands |
| *Cerastium holosteoides* | *Caryophyllaceae* | grasslands, meadows, roadsides |
| *Cerastium semidecandrum* | *Caryophyllaceae* | dry sandy soils, grasslands |
| *Chaetosciadium trichospermum* | *Apiaceae* | dry grasslands, rocky habitats |
| *Chrysanthemum coronarium* | *Asteraceae* | fields, roadsides, coastal habitats |
| *Cichorium endivia* | *Asteraceae* | fields, roadsides, disturbed habitats |
| *Cichorium pumilum* | *Asteraceae* | dry grasslands, roadsides |
| *Cladanthus mixtus* | *Asteraceae* | sandy soils, fields, grasslands |
| *Clinopodium acinos* | *Lamiaceae* | dry grasslands, rocky places |
| *Craspedia variabilis* | *Asteraceae* | grasslands, open woodlands |
| *Daucus broteri* | *Apiaceae* | dry grasslands, rocky slopes |
| *Daucus carota* | *Apiaceae* | meadows, roadsides, grasslands |
| *Echium judaeum* | *Boraginaceae* | dry fields, rocky areas |
| *Echium plantagineum* | *Boraginaceae* | grasslands, roadsides |
| *Erodium cicutarium* | *Geraniaceae* | dry soils, fields, roadsides |
| *Erodium malacoides* | *Geraniaceae* | fields, coastal areas, disturbed sites |
| *Erophila verna* | *Brassicaceae* | dry soils, rocks, grasslands |
| *Euphorbia maculata* | *Euphorbiaceae* | disturbed areas, roadsides, sandy soils |
| *Euphorbia peplus* | *Euphorbiaceae* | gardens, fields, disturbed habitats |
| *Fagopyrum esculentum* | *Polygonaceae* | cultivated fields, disturbed habitats |
| *Glebionis coronaria* | *Asteraceae* | fields, roadsides, coastal areas |
| *Geranium molle* | *Geraniaceae* | grasslands, roadsides |
| *Goodenia vilmorinae* | *Goodeniaceae* | dry woodlands, grasslands |
| *Heliotropium europaeum* | *Boraginaceae* | dry fields, roadsides |
| *Heliotropium hirsutissimum* | *Boraginaceae* | dry fields, rocky habitats |
| *Heliotropium tenuifolium* | *Boraginaceae* | sandy soils, dry habitats |
| *Hirschfeldia incana* | *Brassicaceae* | fields, roadsides, disturbed areas |
| *Hordeum geniculatum* | *Poaceae* | coastal areas, saline soils |
| *Hordeum glaucum* | *Poaceae* | grasslands, disturbed sites |
| *Jasione montana* | *Campanulaceae* | dry grasslands, sandy soils |
| *Lagurus ovatus* | *Poaceae* | coastal dunes, sandy areas |
| *Lavatera cretica* | *Malvaceae* | coastal habitats, roadsides |
| *Linum bienne* | *Linaceae* | grasslands, roadsides |
| *Lobularia maritima* | *Brassicaceae* | coastal areas, rocky slopes |
| *Lomelosia prolifera* | *Caprifoliaceae* | dry grasslands, rocky sites |
| *Malva neglecta* | *Malvaceae* | fields, roadsides, disturbed habitats |
| *Malva parviflora* | *Malvaceae* | disturbed areas, fields |
| *Papaver carmeli* | *Papaveraceae* | rocky slopes, dry habitats |
| *Papaver dubium* | *Papaveraceae* | arable fields, roadsides |
| *Papaver rhoeas* | *Papaveraceae* | arable fields, disturbed habitats |
| *Petrorhagia prolifera* | *Caryophyllaceae* | dry grasslands, sandy soils |
| *Plantago afra* | *Plantaginaceae* | dry grasslands, roadsides |
| *Poa annua* | *Poaceae* | lawns, roadsides, disturbed habitats |
| *Podolepis canescens* | *Asteraceae* | dry grasslands, open woodlands |
| *Podolepis gracilis* | *Asteraceae* | grasslands, sandy soils |
| *Podotheca gnaphalioides* | *Asteraceae* | dry open habitats, sandy soils |
| *Polycarpon tetraphyllum* | *Caryophyllaceae* | dry soils, roadsides |
| *Portulaca oleracea* | *Portulacaceae* | disturbed areas, fields, sandy soils |
| *Ranunculus scandicinus* | *Ranunculaceae* | wet meadows, grasslands |
| *Reseda alba* | *Resedaceae* | dry fields, roadsides |
| *Reseda lutea* | *Resedaceae* | grasslands, roadsides |
| *Rhodanthe chlorocephala* | *Asteraceae* | dry grasslands, sandy soils |
| *Rhodanthe humboldtiana* | *Asteraceae* | arid areas, open habitats |
| *Rhodanthe manglesii* | *Asteraceae* | grasslands, sandy soils |
| *Rhodanthe margarethae* | *Asteraceae* | dry open habitats |
| *Rhodanthe stricta* | *Asteraceae* | dry grasslands, shrublands |
| *Ricotia lunaria* | *Brassicaceae* | rocky slopes, dry habitats |
| *Rudbeckia hirta* | *Asteraceae* | prairies, grasslands, disturbed areas |
| *Scabiosa prolifera* | *Caprifoliaceae* | dry grasslands, rocky sites |
| *Schoenia cassiniana* | *Asteraceae* | dry grasslands, sandy soils |
| *Schoenia filifolia* | *Asteraceae* | dry grasslands, sandy soils |
| *Schoenia macivorii* | *Asteraceae* | arid areas, open habitats |
| *Scrophularia peregrina* | *Scrophulariaceae* | forests, shaded areas, rocky sites |
| *Sedum sediforme* | *Crassulaceae* | rocky slopes, dry habitats |
| *Senecio vulgare* | *Asteraceae* | disturbed areas, roadsides |
| *Sideritis hyssopifolia* | *Lamiaceae* | rocky mountain habitats |
| *Silene aegyptiaca* | *Caryophyllaceae* | dry grasslands, sandy soils |
| *Silene conica* | *Caryophyllaceae* | sandy soils, grasslands |
| *Silene gallica* | *Caryophyllaceae* | fields, roadsides, disturbed habitats |
| *Sinapis alba* | *Brassicaceae* | fields, roadsides |
| *Swainsona formosa* | *Fabaceae* | arid regions, open grasslands |
| *Solanum nigrum* | *Solanaceae* | fields, gardens, disturbed habitats |
| *Sonchus oleraceus* | *Asteraceae* | fields, roadsides, disturbed areas |
| *Stipa capensis* | *Poaceae* | dry grasslands |
| *Tordylium carmeli* | *Apiaceae* | dry grasslands, rocky areas |
| *Trachymene pilosa* | *Araliaceae* | grasslands, open woodlands |
| *Trachymene coerulea* | *Araliaceae* | dry grasslands, sandy soils |
| *Trifolium arvense* | *Fabaceae* | dry grasslands, sandy soils |
| *Trifolium campestre* | *Fabaceae* | grasslands, fields |
| *Trifolium palaestinum* | *Fabaceae* | dry grasslands, fields |
| *Trifolium stellatum* | *Fabaceae* | dry grasslands, rocky sites |
| *Triticum dicoccoides* | *Poaceae* | rocky slopes, wild grasslands |
| *Viola arvensis* | *Violaceae* | arable fields, roadsides |
| *Viola tricolor* | *Violaceae* | grasslands, fields, disturbed habitats |
| *Xerochrysum bracteatum* | *Asteraceae* | grasslands, open woodlands, sandy soils |

